# Conformational switch of a peptide provides a novel strategy to design peptide loaded porous organic polymer for Pyroptosis pathway mediated cancer therapy

**DOI:** 10.1101/2024.03.05.583621

**Authors:** Snehasis Mishra, Achinta Sannigrahi, Santu Ruidas, Sujan Chatterjee, Kamalesh Roy, Deblina Misra, Barun Kumar Maity, Rabindranath Paul, Krishna Das Saha, Asim Bhaumik, Krishnananda Chattopadhyay

**Author notes:** Contributed equally (Snehasis Mishra, Achinta Sannigrahi, Santu Ruidas, Sujan Chatterjee, Kamalesh Roy). Email addresses of the corresponding authors.

## Abstract

While peptide-based drug development is now extensively explored, this strategy does have limitations, which come from their rapid excretion from the body (or shorter half-life in the body), and vulnerability to protease-mediated degradation. To overcome these limitations, here we introduce a novel strategy for the development of a peptide-based anticancer agent using the conformation switch property of a chameleon sequence stretch, which we derived from a mycobacterium secretory protein, MPT63. Then, we loaded this peptide in a new porous organic polymer (PG-DFC-POP) synthesized using phloroglucinol and acresolderivative via condensation reaction for delivering the peptide selectively to the cancer cells. Employing an ensemble and single molecule approaches, we demonstrate that this peptide undergoes a disordered to alpha-helical conformational transition, which is triggered by a low pH environment inside cancer cells. This adopted alpha-helical conformation results in the formation of proteolysis-resistant oligomers, which show efficient membrane pore-forming activity selectively for negatively charged phospholipid which is accumulated in cancer cell membrane. Our *in vitro* and *in vivo* experimental results demonstrate that the peptide-loaded nanomaterial PG-DFC-POP-PEP 1 exhibits significant cytotoxicity in cancer cells, leading to cell death through the Pyroptosis pathway that is confirmed by monitoring numerous associated events starting from lysosome membrane damage to GSDMD-induced cell membrane demolition. We believe that this novel conformational switch-based drug design strategy has great potential in endogenous environment-responsive cancer therapy and the development of future drug candidates to mitigate cancers.

## INTRODUCTION

Although small molecules-based therapeutics are still the major players in the drug development ecosystems, the use of peptides and proteins are gaining momentum, particularly in the field of cancer therapeutics. Among the protein-based drugs, preference has been given to the monoclonal antibody-based biologics molecules (mAbs)^1^, because of their efficient binding to the target receptors for which they have been designed. In addition to the mAbs, there are efforts to design intelligent proteins/peptides based therapeutics^2^, which can bedelivered selectively to pathological tissues without affecting other parts of the body. Specificity is always an important criterion, enabling the reduction of the side effects. In this context, it is to be mentioned that antimicrobial peptides (AMPs) have been found as new drug molecules for targeting cancer cells. Many AMPs, acting as anti-cancer peptides (ACPs), can cross cell membranes and to kill either bacteria or cancer cells. The specific recognition of tumor cells is facilitated by the presence of the negatively charged PS on their surface, caused by high levels of ROS and hypoxia that modify tumor microenvironment and induce membrane phospholipids dysregulation^4^ One example of these peptides/protein-based molecules is LL-37, which generates caspase-independent calpain-mediated apoptosis in Jurkat cells as well as mitochondrial depolarization and caspase-independent apoptosis in human oral squamous cell carcinoma (OSCC) cells SAS-H1^5^. At the same time, LL-37 does not cause cell death of keratinocyte cell line HaCaT, showing that it can be specific and selective for the treatment of OSCC ^6^. Ranatuerin-2PLx (R2PLx) is another 28-amino acid polypeptide, which has been derived from the skin of pickerel frog (Rana palustris). The anti-cancer activity of R2PLx has been demonstrated by the treatment of prostate cancer cell PC-3 which induced AMP-dependent caspase-3 activity and early cell apoptosis^7^. Mastoparan-C (MP-C), extracted from the venom of the European hornet (Vespa crabro), and its analogues, that have been optimized for stability and membrane permeability by cys-cys cyclization and a N-terminal TAT extension, showed anti-cancer activity against non-small cell lung cancer H157, melanocyte MDA-MB-435S, human prostate carcinoma PC-3, human glioblastoma astrocytoma U251MG and human breast cancer MCF-7 cell lines. Importantly, all the peptides show relatively weak activities against the normal human microvascular endothelial cell line HMEC-1. The GW-H1 (GYNYAKKLANLAKKPANALW) peptide is a novel cationic amphipathic AMP effective against several HCC derived cell lines, including J5, Huh7, and Hep3B. Chen et al. used flow cytometry and western blot analysis to show that GW-H1 anti-tumor activity is based on its ability to induce apoptosis in a dose-dependent manner^9^. AMPs as anticancer agents are not without limitations, which included poor stability, proteolytic degradation, potential toxicity, low bioavailability and limited specific delivery to tumor cell^10^. Some of theseproblems can be overcome by different strategies, including chemical modifications, amino acids substitution, fusion with cell penetrating peptides, and nanoparticles loading. It should be mentioned that cancer cell lysosome membrane targeting peptides suffer from instability before reaching out the endolysome membrane due to proteolytic cleavage by different proteases present inside endolysosome. Therefore, it is utterly important to design a peptide that can retain its activity and structure before its interaction with endosomal inner membrane. We hypothesize that the combination of three strategies can help us design a successful peptide-based drug candidates against cancer. First, targetingthose cells by multi phenolic mesoporous nanomaterials (here POP) which can selectively release drug based on environment pH, second, inside mesoporous nanomaterials we can efficiently load a peptide that can form pH responsive oligomers which can be toxic but resistant against proteolytic degradation, third, those oligomers have selective pore forming activity towards negatively charged lipid (i.e PS) which is abundant in cancer cells.

As proof of the concept of the above hypothesis, we introduce here a stimuli sensitive peptide loaded porous organic polymers as potential anticancer agents. Our design of stimuli sensitive peptide uses a strategy of novel conformational switch^11^of a protein, MPT63^12, 13^. We have recently demonstrated that mycobacterium tuberculosis derived MPT63 can adopt a helical structurewhile a β sheet under physiological pH of 7.5 and this switch between β-sheet and α-helix can be regulated by the presence of chameleon sequences. We found that at low pH environmentMPT63 can form toxic oligomers which preferably bind with negatively charged lipids to create membrane pores^14^.Chameleon sequences (ChSeq) are sequence strings of identical amino acids, which can adopt different conformations in protein structures under different conditions. In MPT63, we found eight stretches of chameleon sequences. These are 19VGQVVL24, 27KVSDLK32, 28VSDLKS33, 34SDLKSST40, 56AIRGSV61, 52ATVNAI57, 51TATVNA56, and 69NARTAD74(Figure 1A)^13^. Interestingly, there are two short overlapping stretches of chameleon sequences KVSDLK (residues 27–32) and VSDLKS (residues 28– 33) near the N-terminus. Using refolding simulation and the *ab initio* study, we found that they have the propensity to form helices^12^.We have recently shown that the chameleon sequence in MPT63 makes the protein sensitive towards solutionpH^12^. At physiological pH of 7.4 the protein is at its native state of predominantly β-sheet, and it does not bind to cell membrane. In contrast at low pH,we found that MPT63 binds to membrane, forming toxic oligomeric species which can destabilize lipid bilayer through membrane pore formation^14^. Since cancer cells are believed to have low pH, we hypothesize that carefully designed peptide regions of MPT63 containing the chameleon sequences would selectively target cancer calls (with low pH) without affecting the normal cells (at pH 7.5). For this study, we have selectively chosen two chameleon sequence stretches from MPT63, whichare^19^VGQVVLGWKVSDLKSSTAVIPGYC^40^(denoted as PEP1) and^52^ATVNAIRGSVTPAVSQFNARTADGC^74^(denoted as PEP2). We then loaded synthesized PEP1 onto a newly synthesized porous organic polymers (POPs) (PG-DFC-PEG-PEP1) to investigate if it can target cancer cells selectively using a Colorectal Cancer Cell (CRC)line. Organic materials bearing multiple phenolic-OH sites in the polymer network have shown high therapeutic activity as carrier or drug for cancer treatments^15,16^. In this context, porous organic polymers are an interesting class of materials bearing nanoscale porosity and they are synthesized through copolymerization of reactive organic building blocks. To enhance the abundance of phenolic-OH in the polymer framework^17–20^, herein we have chosen phluroglucinol and diformyl cresol as the building blocks for the synthesis of a new polyphenolic material PG-DFC-POP. We are interested in CRC because it is the third most frequent type of cancer and the second leading reason of cancer-assisted deaths worldwide. Between 2012 and 2030, the new patient case histories are believed to increase by 77%, and the majority of this increase would happen in the low and middle income countries (LMICs)^21,22^. There is an urgent need to develop affordable and specific therapeutic strategies against CRC and we believe that our use of this novel conformational switch strategy towards designing new Peptides would help combating not only CRC, but other cancers as well.

**Figure 1:**
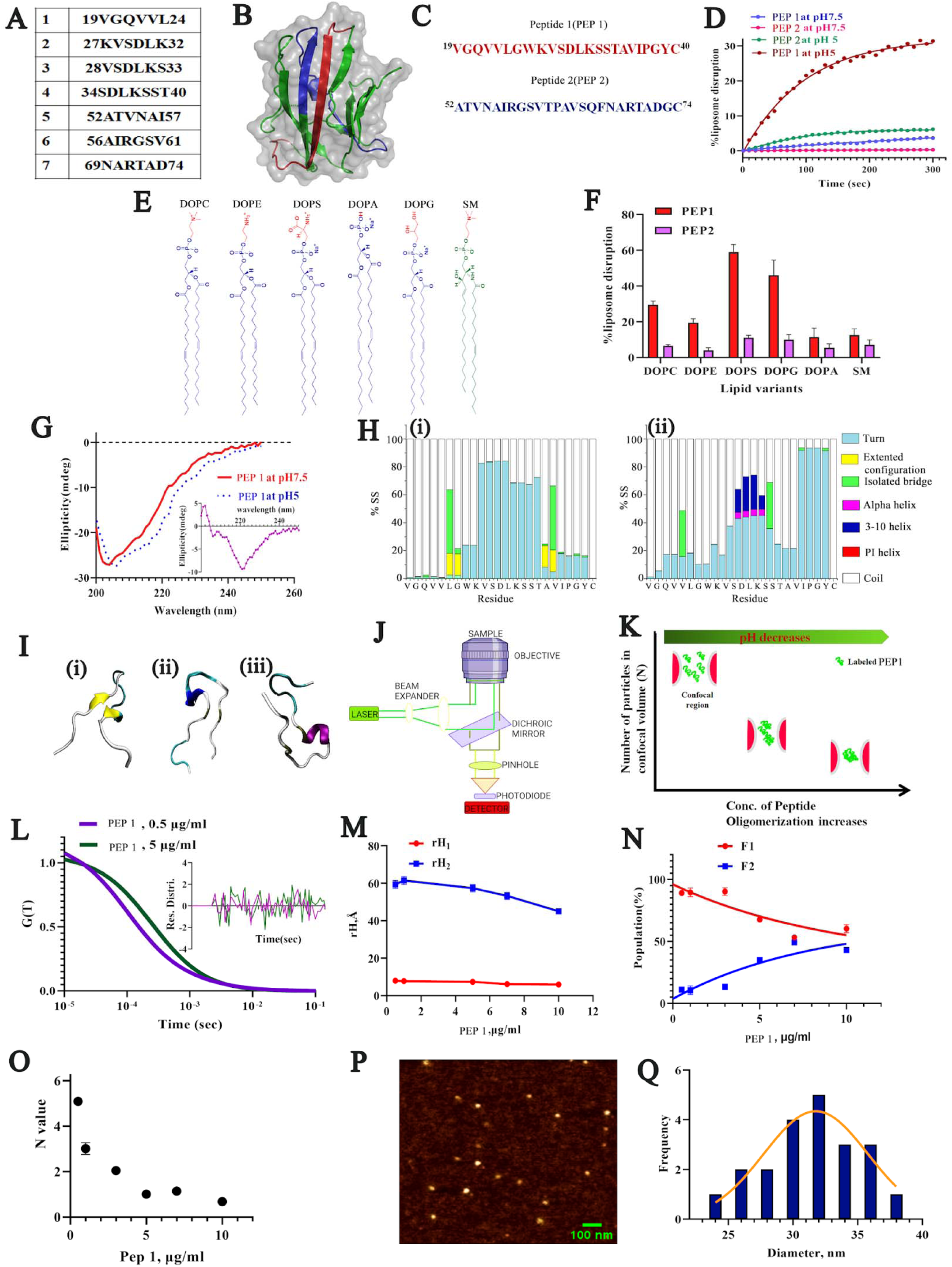
Conformational switch in MPT63 derived peptides: (A). Chameleon stretches in MPT63 determined from ChSeqdatabase.(B). Pymol structure of MPT63 (PDB ID 1LMI). The chameleon sequence (19VGQVVLGWKVSDLKSSTAVIPGYC40(PEP1) a 52ATVNAIRGSVTPAVSQFNARTADGC74(PEP2)) are marked by red and blue respectively. (C). O selected peptide sequences for this study. (D). Leakage experiments of calcein dye entrapped DOPC SUV when treated with PEP1 and PEP 2 at two different pH conditions. (E). Structures of different lipids. (F). PH shows the percentages of calcein dye leakage as pore forming ability of PEP 1 and PEP2 from liposom composed of different lipid components. (G). Secondary structure data of the peptides show the conformational switch in PEP 1 from unstructured and beta sheet conformation to alpha helical conformational while the solution pH changes from 7.5 to 5. Inset shows the subtracted CD data (CD data at pH 5-CD data pH 7.5) suggesting the existence of alpha helical signature (222 nm) in PEP 1 at pH 5. (H). Secondary structural propensity percentage of PEP1 at pH (i) 7.5 and (ii) pH 2 as obtained from MD simulation analyse (Extended configuration stands for parallel and antiparallel beta sheets). (I). Secondary structure of PEP1 pH (i) 7.5 andboth [(ii),(iii)]at2 as obtained from MD simulation production run. (J). A typical setup of FC instrument. (K). The representative plot of the number of Alexa 488 maleimide labeled PEP 1 molecules confocal volume against the concentration of PEP1. (L) FCS correlations were fitted using two component gaussian function. These correlation data were measured for PEP 1 at pH 5. Inset plots indicate the residue distribution which suggests the goodness of the fit. (M).Hydrodynamic radii of the fast component (rH1) at slow component (rH2) of PEP 1 at pH 5 were plotted against the increasing concentration of the PEP 1. (N) With increasing concentration of PEP 1 at pH 5, monomeric component (F1) decreases and simultaneously increase in oligomeric component (F2) was found. (O). The number of particles of PEP 1 in the conformational volume at pH 5 with increasing concentration of PEP1. (P). AFM topographic image of PEP1 at pH 5. The scale bar for the image is 100 nm. (Q). Histogram of the particle size distribution of PEP 1 oligomers at pH as estimated from the AFM image using Fiji Image J software.

The *in vitro* and *in vivo* experiments presented here show that PG DFC POP PEP1 exhibits significant cytotoxicity in cancer cells, leading to cell death through the Pyroptosis pathway that is confirmed by monitoring numerous associated events starting from lysosome membrane damage to GSDMD-induced cell membrane demolition.To the best of our knowledge, this is the first report of designing protein-based cancer therapeutic using pH sensitive conformational switch-based strategy.

## RESULTS AND DISCUSSION

### PEP1 (but not PEP2) is sensitive to low pH-induced conformational switch leading to membrane pore formation

MPT63 is a secretory immunogenic protein (17 kDa, PDB ID 1LMI) of mycobacterium tuberculosis with an overallβ**-**sandwich fold (Figure 1B). We have recently shown that at low pH condition (pH 5), MPT63 acquires significant alpha helical confirmation^12,14^. Our results indicated that the sequence region 19-30 is responsible for the pH responsive conformational switch event of MPT63, which leads to the formation of toxic oligomers of MPT63. These toxic oligomers disrupt the cellular membrane through pore formation.

Ch. Seq. data analysis shows the presence of a number of chameleon sequence stretches in MPT63 (Figure 1 A). Here, we have selectively chosen two chameleon sequence stretches, namely^19^VGQVVLGWKVSDLKSSTAVIPGYC^40^(PEP1)and^52^ATVNAIRGSVTPAVSQFNAR TADGC (PEP2) for this study (Figure 1B,C). We modified the peptide sequences with cysteine residue to increase the oligomerisation propensity through disulfide linkage. The hydrophobic contribution of each amino acid constituting PEP1 and PEP2 were analyzed using ProtScale (http://web.expasy.org/protscale/) (Figure S1A). We used the hydrophobicity scale as defined by Kyte and Doolittle. Figure S1A shows higher hydrophobicity for PEP1 in comparison to PEP2.

We then used Aggrescan analysis (http://bioinf.uab.es/aggrescan/), which predicted higher oligomerisation propensity for PEP1 when compared to PEP2 (Figure S1B). Since, low pH induced conformational change and oligomerization leading to the membrane damage through pore formation is the objective of this study, we first examined the membrane pore forming activity of PEP1 and PEP2 under two pH conditions by measuring calcein release from synthetic DOPC liposomes (Figure 1D). At pH 7.5, neither PEP1 nor PEP2 showed any significant calcein dye leakage from the dye entrapped liposomes (Figure 1D). However, when we carried out this assay at pH 5, we found significantly higher calcein release for PEP1 (30%) when compared to PEP 2 (6%) (Figure 1D). It is well known that cancer cells possess high surface expression of negatively charged lipid components PS, and hence we subsequently carried out the pore formation experiment using different lipids based on their head group variations (Figure 1E). We found significantly higher leakage in negatively charged lipid vesicles by PEP1 at pH 5 condition (Figure 1F). In contrast, PEP2 did not show pore formation(Figure 1F). Based on this observation we considered PEP1 to be a better molecule to develop potential pore-forming anticancer agent, which has higher sensitivity towards low pH condition prevalent inside cancer cells. Next, to understand the conformational property of PEP1, we employed far UV CD spectroscopy to show that PEP1 ismostly disordered at pH 7.5, although a small shoulder at 216nm was observed which indicates the presence of small extent of β-sheet conformation (Figure 1G). In contrast, when the peptide was subjected to pH 5, we observed an increase in the ellipticity at around 220-222nm (Figure 1G). This increase takes place due to an increase in the helical content of PEP1 at pH 5(Figure 1G,inset).Computational analyses using molecular dynamic simulations with PEP1 provided complimentary information on higher alpha helical propensity at low pH (Figure 1Hi,ii,Figure 1I and Figure S1C,D).In order to verify the conformational changes within the chameleon sequence stretches(PEP1) present in MPT63, classical atomistic molecular dynamics simulations have been performed with the help of the NAMD program^23^. Initially, to mimic the experimental conditions, two (pH7 and pH2) input models of PEP1 have been constructed. For the purpose of verification of random coil to helix transformation, protonation states of the titratable residues (VAL, LYS, ASP, TYR, CYS) were changed based on the pKa/pH correlation of the amino acids via CHARMM-GUI input^24^ generator. Atomistic molecular dynamics reveals that, at pH 7 condition beta strand is preferred between ^6^LEU-GLY^7^and ^18^ALA-VAL^19^ and there is no significant propensity towards helix formation Figure 1Hi. In contrast, at low pH, ^12^ASP-LEU-LYS-SER^15^stretch showed 3/10 helix as well as alpha helical propensity (Figure 1Hii,I).

The pore formation by proteins and peptides has been extensively studied. It is believed that pore formation occurs either through monomeric protein or through self-association. To determine if PEP1 self-associates, we used Fluorescence Correlation Spectroscopy (FCS). FCS is a popular spectroscopic method (Figure 1J) to monitor molecular diffusion of a labeled molecule at single-molecule resolution^25^. An optimum fit of FCS data would provide an estimation of hydrodynamic radius (r_H_) and the number of particles in the confocal volume (N). As shown in Figure 1K, a self-associating Peptide system would increase r_H_and decrease N. We performed FCS experiments using 0.5 µg/ml Alexa488-labeled PEP1, which we mixed with increasing concentrations of the unlabeled PEP1 (but keeping the concentration of the labeled Peptide constant at 0.5 µg/ml). Since increase in concentration would favor self-association, using this experiment we could monitor self-association of the peptide using increasing effective concentration.With PEP1 at pH5 (the helical form), we found a shift toward right (representative data for the PEP1 at pH 5 are shown in Figure 1K,L). We fit the correlation functions of the PEP 1 using a model of two components, in which the fast-diffusing component (r_H1_) was the monomer, while the slow component (r_H2_) represented a high-molecular-weight species, or oligomers. The r_H_ value calculated for the fast component was found to be ∼8.5 Å (Figure 1M). In contrast, the slow component showed a r_H_ of ∼60 Å (Figure 1M). Interestingly, at pH 5 with increasing concentration of PEP1, the amplitude of the monomer component (F1) decreased, while that of the slow high-molecular-weight oligomer component (F2) increased (Figure 1M).

The number of particles (N) decreased with increasing concentration, which suggested that PEP1 self-associated to form high molecular-weight oligomers at pH 5 (Figure 1M,N,O). We then employed AFM for direct visualization of the oligomers (Figure 1P). As expected, we did not find any large molecular species with PEP1 at pH 7.5(Figure S1E). In contrast, PEP1 at pH5 formed some round shaped oligomeric particles with 90% population of average size of 30 nm in diameter (Figure 1P,Q). In this context, it should be mentioned that proteolytic degradation of the peptide based anticancer agent is the principal drawback behind the drug development strategy^26^. Therefore, to explore the peptide susceptibility to proteolysis, PEP1 was incubated at pH5 with trypsin at 1:200 enzyme :substrate weight ratio at 24 °C, and then analyzed by FCS, which showed similar correlation function as observed for untreated control (Figure S2A). Our FCS data analysis suggested almost equal populations and size of the oligomers for both treated and untreated condition (Figure S2B). This data indicates that PEP 1 oligomers formed at pH 5 seem to be resistant towards proteolytic degradation.

### Synthesis of PG-DFC-POP

PG-DFC-POP was prepared via solvothermal synthesis^27^strategy using phloroglucinol and diformylcresol as starting materials (Figure 2). Powder X-ray diffraction analysis was carried out to understand the structural property of PG-DFC-POP. Figure 3A shows a broad peak at 10-25° 2Θ which reveals amorphous nature of the POP. The permanent porosity and specific surface area of PG-DFC-POP was assessed from N_2_ adsorption– desorption analysis at 77 K. As shown in Figure 3B adsorption isotherm reveals a large volume of nitrogen uptake that has occurred at low relative pressure region resulting a type I isotherm. This indicates the microporous nature of PG-DFC-POP. The estimated (Brunauer-Emmett-Teller) surface area of PG-DFC-POPwas 335 m^2^ g^_1^with total pore volume of 0.019 cc g^-1^. Furthermore, the pore size distribution (Figure 3B, inset) was obtained from nonlocal density functional theory (NLDFT) which implies the presence of micropores with pore dimension of 1.6 nm in PG-DFC-POP. The structural growth and bond connectivity of PG-DFC-POP was determined by FT-IR (Figure 3C) where, it was found that a broad peak at 3389 cm^-1^that indicates the presence of terminal OH functionality throughout the material and a peak at 2900 cm^-1^ can be assigned for the stretching of alkyl that indicates the formation of PG-DFC-POP by the reaction of phloroglucinol and diformylcresol by the elimination of water. After the incorporation of peptide in PG-DFC-POP a stretching frequency at 3329 cm^-1^ was appeared due to hydroxyl group of carboxylic acid moiety and –NH functionality of amine moiety from the peptide part and further signal at 1644 cm^-1^ is due to carbonyl functionality of carboxylic acid. Further, C solid NMR (Figure S3) spectrum shows a peak at 58 ppm corresponding to the connecting aliphatic carbon containing hydroxyl group. On the other hand, peaks at 21 and 150 ppm could be attributed to methyl carbons and carbon atomsof the aromatic benzene rings, respectively. A small peak at 180ppm can be assigned for keto carbon.To understand the morphological features of PG-DFC-POP transmission electron microscopy (Figure 3Da and b) and scanning electron microscopy (Figure 3D c and d) images were used, which revealnear spherical morphology with particle dimension of 0.5-2.50 µm. The HR TEM images of PG-DFC-POP-PEP before and after the loading of the peptide suggested no change in particle morphology of the POP (Figure 3D a and b).

**Figure 2:**
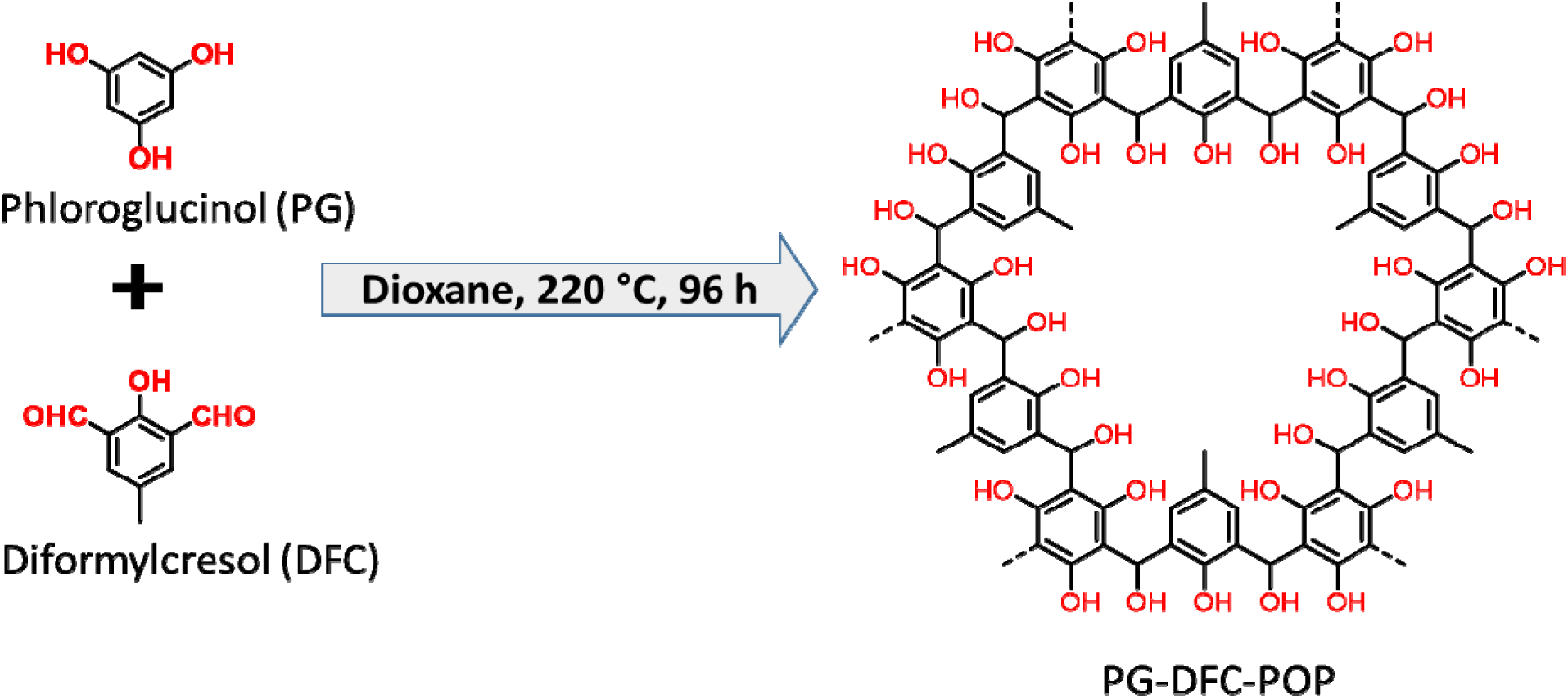
Synthesis strategy of PG-GFC-POP using phloroglucinol and diformylcresol.

**Figure 3:**
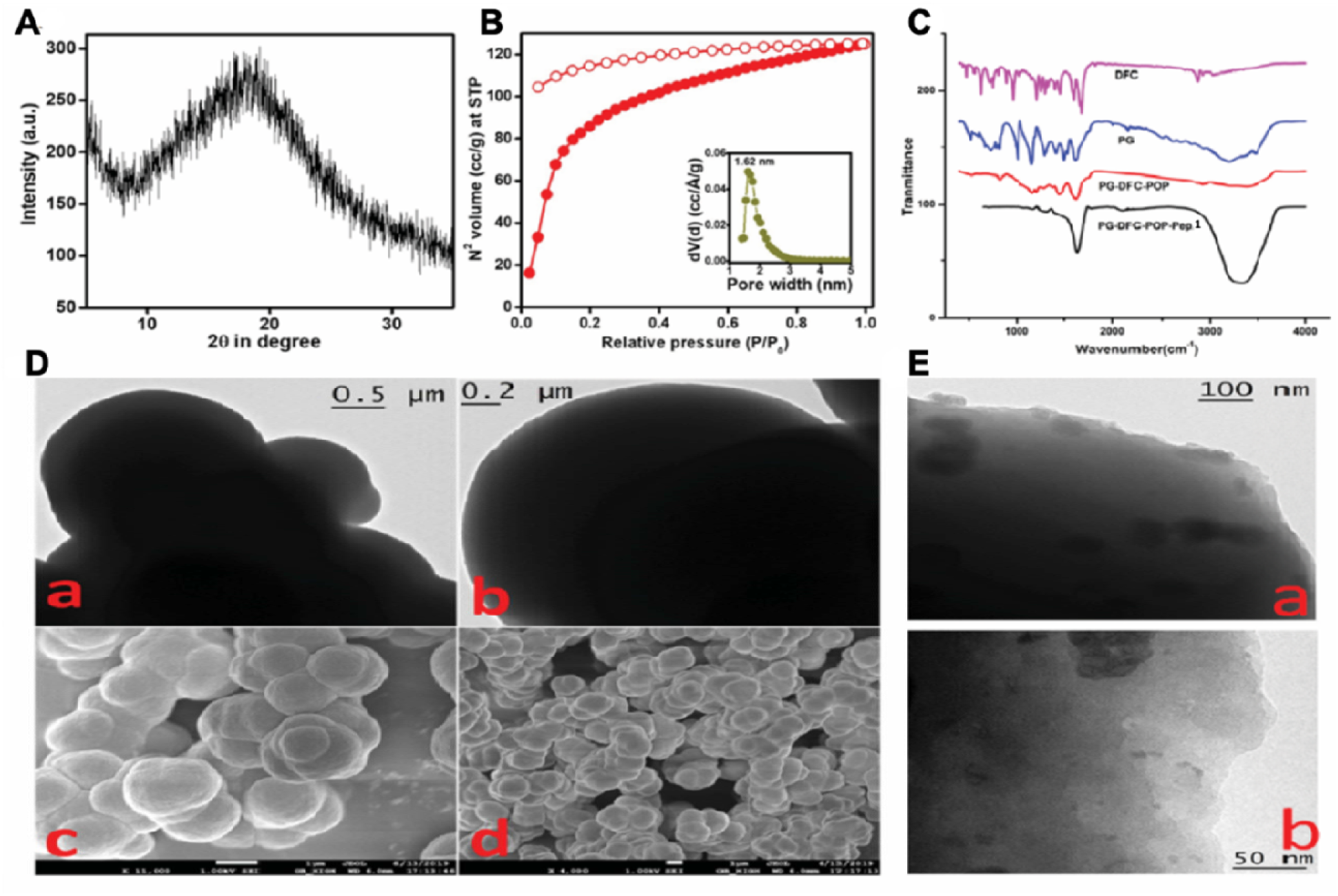
(A) Powder X-ray diffraction pattern of our synthesized PG DFC POP, (B) N_2_ adsorption desorption isotherm of PG-DFC-POP and respective NLDFT pore size distribution plot as shown in inset (C) FTIR spectral map of PG-DFC-POP, PG, and DFC. (D) (a, b) HR-TEM and (c, d) FE-S**EM** images PG-DFC-POP. (E) (a, b) TEM images of PEP1-loaded PG-DFC-POP.

### Loading Efficiency of PEP1 in PG-DFC-POPand solution pH dependent release

18 mg of PG-DFC-POP was dissolved in 1 mL of deionised water containing 12 mg of PEP1, and the mixture was stirred for 24 h. After centrifugation, the PG-DFC-POP PEP 1 was obtained as a precipitate. UV measurement determined the concentration of the unloaded PEP 1 (in solution), allowing for the calculation of its weight. The loading efficiency of PEP1 in PG-DFC-POP was determined to be 9.5%.Subsequently, we studied the temporal release of PEP1 from PG-DFC-POP PEP 1 by incubating the conjugated nanomaterials in three different pH solutions and then separating the released PEP1 fraction from the loaded fractions through analytical ultracentrifugation(100,000 g). Our results showed significantly higher released rate and extent of PEP1 from PG-DFC-POP PEP 1 at pH5 (the pH inside endolysosome) (Figure S4).

### *In vitro* experiments against colon cancer cell line suggested dose dependent cell death by PG-DFC-POP-PEP1treatment

As described before, we used PG-DFC-POP-PEP1 (PEP1 loaded PG-DFC-POP) as the active material.The presence of multiple phenolic groups within the framework of this POP significantly enhances its affinity for cancer cells. Consequently, when PEP1-loaded POP materials are employed, they exhibit a pronounced selectivity in transporting the Peptides specifically to cancer cells. This selective transport mechanism leverages the inherent structure of the POP, as its phenolic groups facilitate a strong attraction towards cancer cells, thereby aiding in targeted drug delivery to combat the disease. Since HCT-116 shows several functions like primary as well as secondary metastatic colon cancer, we used this as a model to determine *in vitro* optimum therapeutic effective dose of PG-DFC-POP-PEP 1 against colon cancer. We found 50% cell death after 5µg/ml of PG-DFC-POP-PEP1 treatment. Interestingly, when we used PEP1alone (in the absence of PG-DFC-POP loading) or only PG-DFC-POP (in the absence of PEP1), we observed significantly less efficacy in restricting cell survivability (p<0.1). Since the administration of dose condition higher than 5µg/ml PG-DFC-POP-PEP1did not yield significantly improved outcome, we considered 5µg/ml as our optimum effective dose for further experimental studies (Figure4A). Morphometric analysis illustrated a significant alteration in cell morphology with effective apoptosis denoting blebbing with 5µg/ml PG-DFC-POP-PEP 1 treatment for 4hr and 8hr. Analysis also asserted notably better remedial efficacy by depicting morphological features of apoptosis-like plasma membrane blebbing (denoted by black arrows), formation of apoptosome (denoted by blue arrows) in HCT-116 colon cancer cells after 5µg/ml of PG-DFC-POP-PEP1 treatment (p<0.01) (Figure 4B).

**Figure 4:**
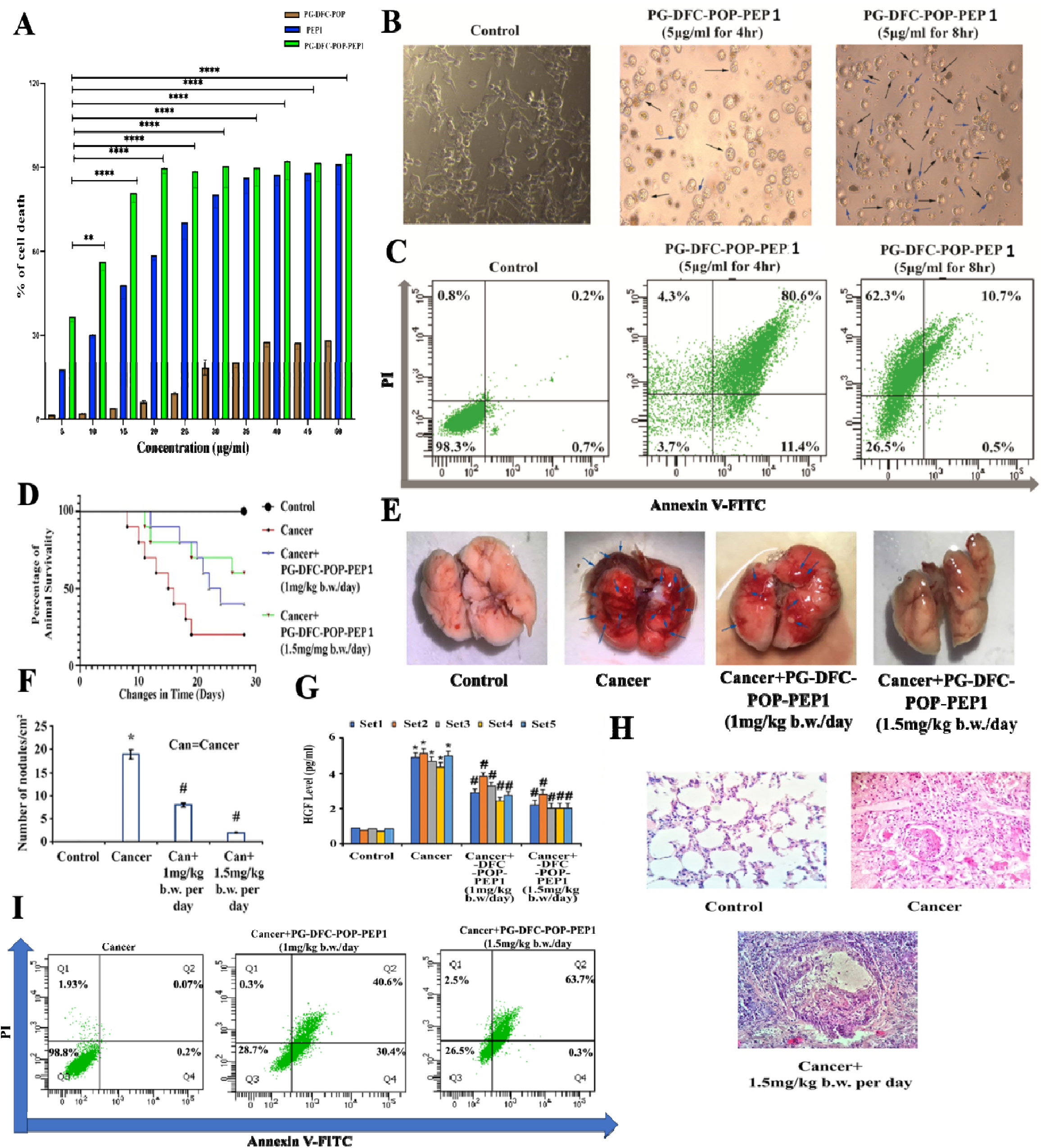
(A-C) Investigation of Cellular Toxicity and Morphological Changes Induced by PG-DFC-POP PEP1 Treatment in HCT-116 Cells: (A) Assessment of Cellular Viability: We performed MTT assay evaluate cellular viability after exposure to PG-DFC-POP, PEP1, and PG-DFC-POP-PEP1. Morphological Alterations: Bright field microscopy images captured at various time points (4 and 8 hours following PG-DFC-POP-PEP 1 treatment (5µg/mL) depict notable changes in cellular morphology( Apoptosis Analysis: AnnexinV-FITC/PI data presentation illustrates apoptotic events in HCT-116 cells up PG-DFC-POP-PEP1 treatment in a time-dependent manner (4 and 8 hours).(D-I) *In vivo* Evaluation of Therapeutic Potential of PG-DFC-POP-PEP1:(D) Survival Analysis: Kaplan-Meier analysis tracks the 30-d survival of CT-26 tumor-bearing mice post-administration of PG-DFC-POP-PEP 1 (1 and 1.5 mg b.w/day).(E) Metastatic Progression: Assessment of metastatic changes following PG-DFC-POP-PE treatment.(F) Nodule Density: Quantification of the number of nodules per square centimeter before and after PG-DFC-POP-PEP1 treatment (1 and 1.5 mg/kg b.w/day).(G) HGF Levels: Evaluation of HGF levels before and after PG-DFC-POP-PEP 1 treatment (1 and 1.5 mg/kg b.w/day).(H) Histological Examination Hematoxylin and Eosin (H & E) staining of *in vivo* tumor tissues post PG-DFC-POP-PEP1 treatment.(I) *vivo* Apoptosis Analysis: Presentation of AnnexinV-FITC/PI data from single cells isolated from *in vivo* samples

We have also assessed PG-DFC-POP -PEP 1’s cytotoxicity in CCD 841CoN and HEK293 cell lines, noting a considerably higher IC50 value compared to HCT116 (Figure S5). This suggests that PG-DFC-POP-PEP 1 exhibits target specificity towards colon carcinoma.We then used the morphology of AnnexinV-FITC+ HCT-116 cells as a marker of apoptosis^29^. From these experiments, we found that the treatment of 5µg/ml PG-DFC-POP-PEP1 increased cell death in a time dependent manner (Figure 4C). We observed significant increase in apoptotic cell population in treated groups suggesting the possible application of PG-DFC-POP-PEP1as a treatment against colon cancer. Since 4h of 5µg/ml drug treatment demonstrated notable morphological features of cellular apoptosis, we selected this dose and time as the lowest effective (optimized) dose for further *in vitro* experimental analysis.

### Experiments in cancer cell induced metastatic tumor model demonstrated the inhibition of tumor metastasis

For determination of *in vivo* effective dose of PG-DFC-POP-PEP1, we performed Kaplan-Meier survivability analysis (Figure4D). We observed significant reduction of tumor metastasis in the untreated group (denoted as Cancer from now onwards, Figure4E) of mice after58 days of experimental tenure (28days of tumor development +30days after dosing). One month post treatment with intravenous PG-DFC-POP-PEP 1 treatment yields significant enhancement in animal survivability rate in a dose-dependent manner. The number of nodules was also calculated (Figure 4F), and it shows a significant decrease after PG-DFC-POP-PEP1 treatment at 1 and 1.5mg/kg bodyweight treatment.

We thencompared the level of the Hepatocyte Growth Factor (HGF) as a marker for metastasis cell progression between the treated and untreated groups. We found elevated HGF level in animals of untreated group, which reduced significantly upon PG-DFC-POP-PEP 1 treatment (Figure 4G). We complemented this by studying the morphology, which indicated significant dose-inducedmitigation in affected foci (marked as blue arrows) area and reduction in average number of nodules in treated groups as compared with untreated animals (Figure 4E,4F).

The comparison involved performing HE staining on the target organ from both the untreated and treated groups, each matched with their respective control samples. The histopathological examination revealed the presence of numerous inflammatory cell clusters, along with apoptotic bodies, macrophages, and cells with condensed nuclei in the treated sets (Figure 4H). We then used Flowcytometric analysis after apoptotic marker AnnexinV-PI staining, which demonstrated a significantand dose dependent increase in apoptotic cell population upon PG-DFC-POP - PEP1treatment (Figure 4I) confirming that apoptosis is the prime reason for the suppression of metastatic nodules at lung and the reduction of HGF level in CT26 mediated pulmonary metastatic tumor model. This in turn not only induced secondary tumor regression but also demonstrated a potential treatment strategy against colon cancer. For the mechanistic studies demonstrated below, we used the dose of 1.5mg/kg b.w./day of PG-DFC-POP-PEP1 treatment for 30 days.

### PG-DFC-POP-PEP1 treatment initiated Pyroptosis through increased N-Gasdermin D (N-GSDMD) level

As described above, we observed an increase in the early as well as late apoptotic cell population after PG-DFC-POP-PEP1 treatment in both *in vitro* and *in vivo* studies. Our results also show a notable shift at AnnexinV-FITC+ cell populations upon treatment towards the marginal site of the late-apoptotic and necrotic quadrate (Figure. 4C and 4H). Major shift in affected cell population (81.8% and 73.0% under *in vivo* and *in vitro* conditions respectively) indicated loss of DNA during programmed cell death. We then wanted to understand if the observed lysosomal localization happens through an active N-GSDMD induced apoptosis, which is very important for the elimination of early colonized metastatic cells at normoxic secondary organs. We observed a significant trend of elevation in active N-GSDMD level after 4hr and 8hr of PG-DFC-POP-PEP1 administration in HCT-116 cells (p<0.01). We also observed a noticeable extent of lysosomal trans-localization of active N-GSDMD (Figure 5A, 5B and 5C) with the Mean fluorescence Intensity (MFI) calculation of Alexa 488 conjugated PEP1. *In vivo* experiments show significant increase in N-GSDMD level (Figure 5D) and its localization as a result ofPG-DFC-POP-PEP1 treatment in a CT26 induced pulmonary metastatic colon cancer model (p<0.01). The MFI calculation further confirms a notable increase in N-GSDMD levels correlating with PEP1 internalization (Figure5E, 5F). To summarize, a significant increase at the level and lysosomal distribution of N terminally cleaved active isoform of Gasdermin confirmed initiation of lysosome mediated programmed cell death pathways, which were induced by our novel Peptide-conjugated material.

**Figure 5:**
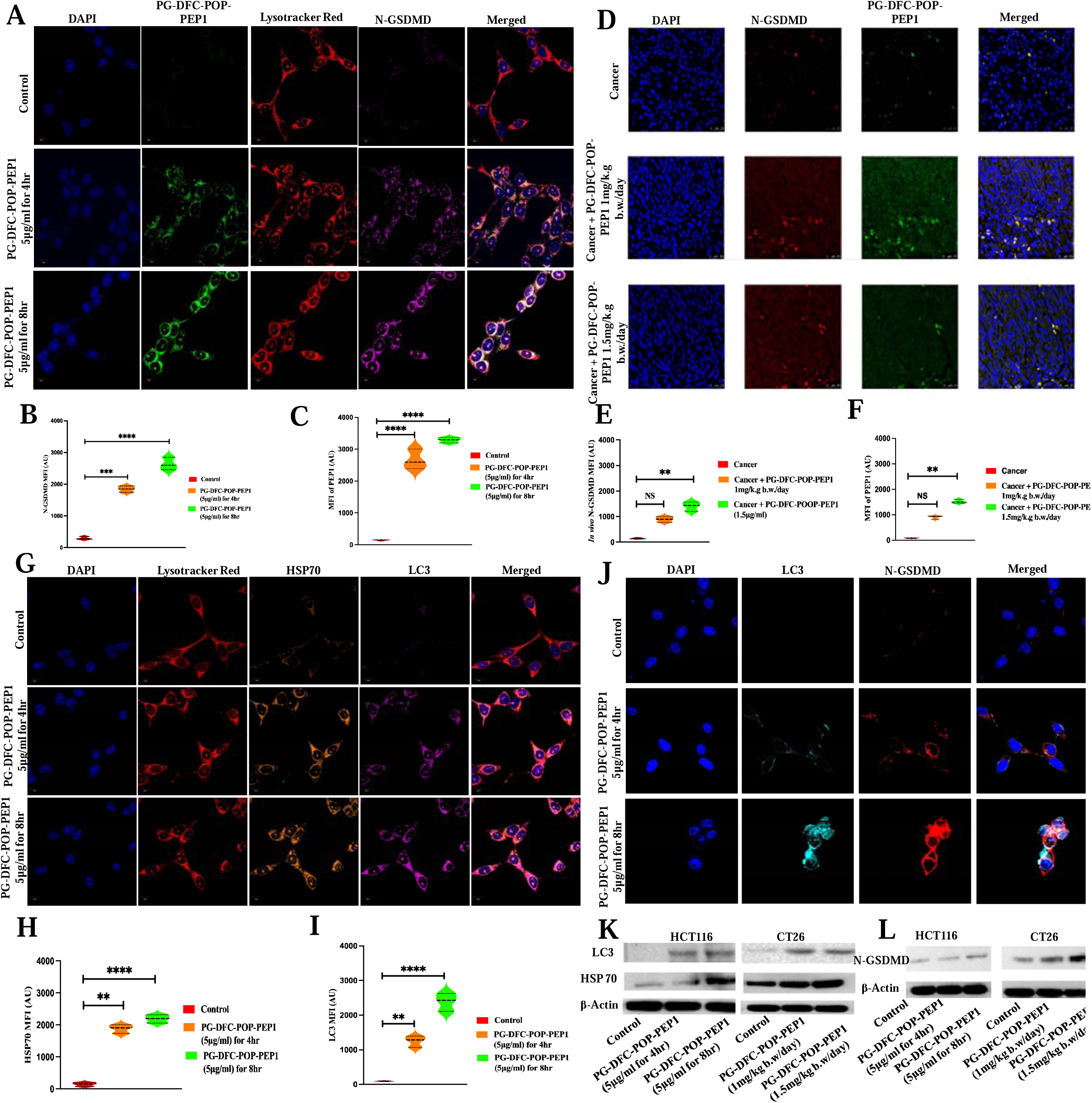
In this figure, we present a comprehensive analysis of the effects of PG-DFC-POP-PEP treatment both *in vitro* and *in vivo.* (A) Photomicrographs depict the expression and localization of the Pyroptosis marker N-GSDMD, visualized using PEP1 tagged with Alexa 488. To highlight cellular components, Lysotracker Red and DAPI were utilized for staining Lysosomes and Nuclei, respectively. (B & C) MFI calculation of N-GSDMD and Alexa 488 conjugated PEP1 from (A) with corresponding statistical analysis. (D) Immunohistochemistry was performed to assess N-GSMD expression, and internalization of Alexa 488 conjugated PEP1 was quantified. (E & F) MFI calculation of N-GSDMD and Alexa 488 conjugated PEP1 from (D) with statistical analysis. (G) Examination of LC3 and HSP70 presence in HCT116 cells post PG-DFC-POP-PEP 1 treatment, with Lysotracker Red and DAPI staining aiding in: cellular context visualization. (H & I) MFI calculation of HSP70 and LC3 from (G) with statistical analysis (J) A closer examination of the crosstalk between LC3 and N-GSDMD, magnified for better visualization both with and without PG-DFC-POP-PEP 1 treatment in HCT116 cells. (K-L) Western Blotting analysing showing estimation of LC3 and HSP70 protein expression post PG-DFC-POP-PEP1 treatment, both *in vitro* and *in vivo.* For *in vitro* studies, HCT-116 and CT-26 cell lines were treated with 5µg/mL PG-DGC-POP-Pep1 for 4 and 8 hours, while *in vivo* experiments involved treatment with 1 and 1.5mg/kg b.w/day of PG-DFC-POP-PEP1. This comprehensive approach allowed assessment of treatment effects under physiological conditions. Statistical comparisons were made using ANOVA test for equality of medians, with **, ***, and **** indicating significant differences (p < 0.05, p < 0.005, and p < 0.001, respectively), while NS denote nonsignificant results. Data are presented as mean ± SEM.

### Alterations in regulatory factors associated with N-GSDMD mediated programmed cell death after PG-DFC-POP-PEP1treatment

It has been reported that externalization of phosphatidylserine is dependent upon lysosomal localization of N-GSDMD^32^.Recent studies have also suggested that the activation of Hsp70-LC3 axis followed by their lysosomal translocation induces Pyroptosis mediated programmed cell death^33^. To determine if the lysosomal localization of N-GSDMD observed here can also lead to activation of Hsp70-LC3 axis, we used confocal microscopy. We found the co-localization of Hsp70 and LC3 proteins within lysosome upon 5µg/ml PG-DFC-POP-PEP 1 treatment for 4hr and 8h (Figure 5G), along with the MFI calculation (Figure 5H, 5I). Using a separate experiment, we have also shown that the treatment results in the expression of N-GSDMD and LC3. The present data also indicated significant elevation in co-distribution of Pyroptosis regulatory LC3 along with N-GSDMD (Figure 5J) in treated *in vitro* sets. Immunoblot analysis from lysosomal fraction similarly exhibited increased LC3 and Hsp70 levels after scheduled PG-DFC-POP-PEP1 treatment in both *in vivo* and *in vitro* sets (Figure 5K and 5L). To summarize, we found that PG-DFC-POP-PEP1 treatment leads to Hsp70-mediated lysosomal translocation of LC3 augmented N-GSDMD activity, resulting in the decrease in colon cancer cell populations. *In vivo* validation confirmed similar increase in co-distribution of LC3 along with HSP70 in 1.5mg/kg b.w./day DFC-POP-PEP1 treated sets as compared with untreated mice (p<0.01) (Figure 5F).

### PG-DFC-POP-PEP1 mediated inflammasome formation in colon cancer

As shown above using *in vitro* and *in vivo* studies, we demonstrated significant increase in proinflammatory LC3, which turned on N-GSDMD mediated programmed cell death in colon cancer cells as well as for *in vivo* metastatic models. This finding prompted us to analyze the status of inflammatory cytokines and related regulatory proteins (Figure 6A-6J). Using confocal microscopy analysis, we find significant increase in IL1β and NLRP3 levels in both *in vitro* and *in vivo* treated groups. We also find an increase in caspase1 and Cathepsin B expression along with their co-distribution with NLRP3 and IL1β respectively. Co-distribution of these proteins and their colocalization in *in vivo* analysis indicated that PG-DFC-POP-PEP1 treatment induces inflammasome mediated apoptotic pathways in cancer cells through uplifting IL 1β mediated NLRP3-LC3-NGSDMD axis that finally induces Cathepsin B and Caspase 1 activity in treated groups. This in turn induces programmed cell death in HCT-116 colon cancer cell line as well as CT26 induced pulmonary metastatic tumors. HE staining of target organ from untreated and treated groups was compared with respective controls. Histopathological observations established numerous inflammatory cell clusters along with apoptotic bodies, macrophages, and pyknotic nuclear-containing cells in treated sets. This confirmed that the treatment induced inflammasome-mediated apoptosis in PG-DFC-POP-PEP1-treated mice.

**Figure 6:**
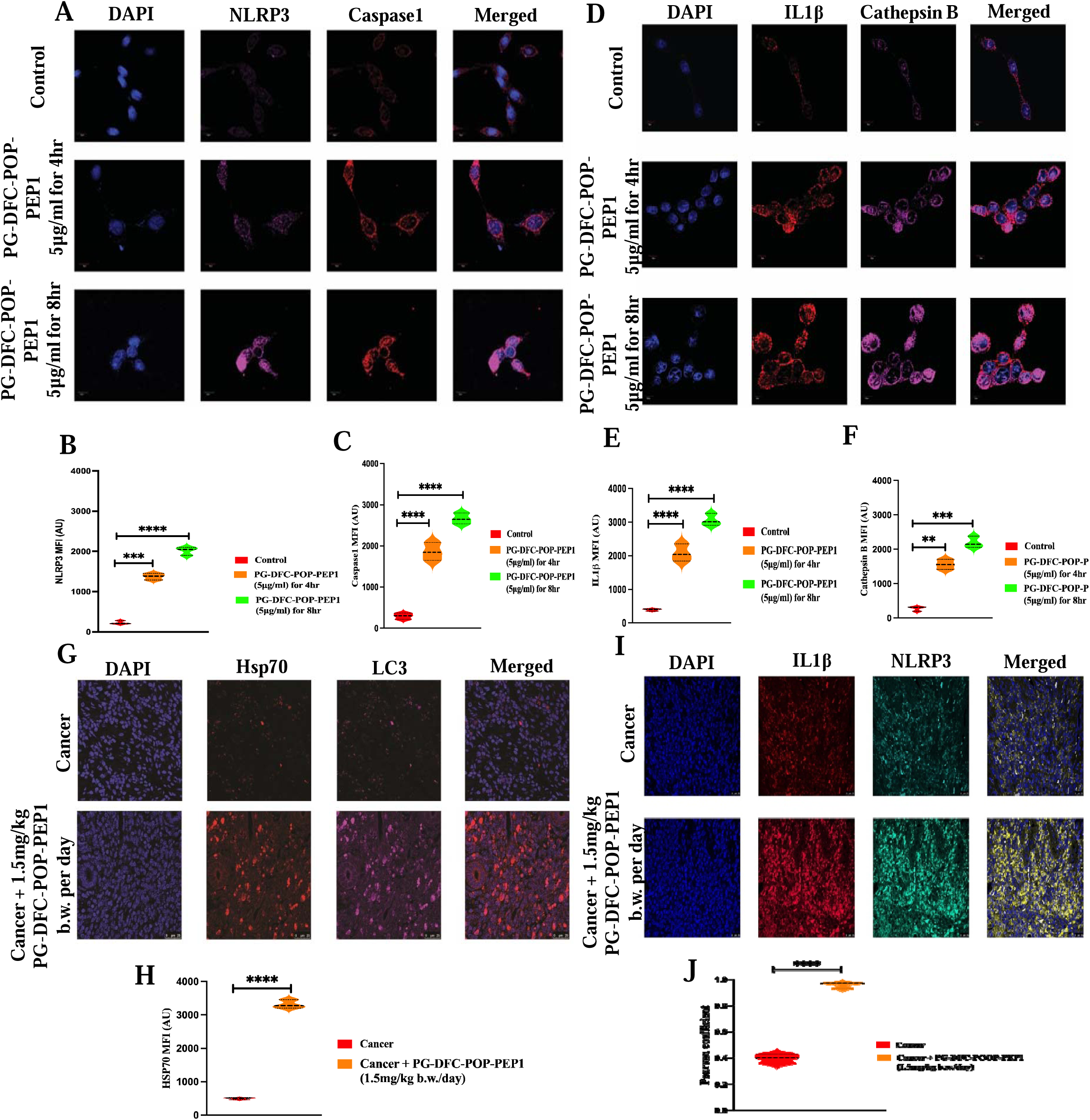
(A-C) Analysis of NLRP3 and Caspase1 Expression: The expression patterns of NLRP3 and Caspase 1, key players in the NLRP3-inflammasome activation pathway. The aim is to elucidate how these components are influenced by PG-DFC-POP-PEP1 treatment (5µg/mL) in a time-dependent manner. The dynamic changes in NLRP3 and Caspase 1 expression are meticulously examined using advanced Confocal Microscopy techniques. (B & C) MFI calculation of NLRP3 and Caspase1 from (A) with corresponding statistical analysis. (D) Assessment of IL-1β and Cathepsin B: The investigation further extends to the expression profiles of IL-1β and Cathepsin B before and after PG-DFC-POP-PEP1 treatment. Understanding how the levels of these critical mediators evolve in response to treatment is pivotal in unraveling the intricacies of NLRP3-inflammasome activation. High-resolution Confocal Microscopy enables precise visualization and quantification of these molecules, shedding light on the mechanistic interplay. (E & F) MFI calculation of IL-1β and Cathepsin B from (D) with corresponding statistical analysis between the groups. (G & H) Comprehensive evaluation of inflammasome markers HSP70 and LC3 expression in an *in vivo* setting following PG-DFC-POP-PEP1 treatment with the statistical analysis. (I) Assessment of IL1β and NLRP3 Colocalization *in vivo:* The investigation extends into the *in vivo* context, scrutinizing the expression levels of two pivotal inflammasome markers, IL1β and NLRP3, both before and after treatment with PG-DFC-POP-PEP 1 at a dosage of 1.5mg/kg b.w per day. Understanding how these markers evolve in response to treatment is crucial in deciphering the intricate molecular dynamics underlying inflammasome activation. This analysis provides valuable insights into the potential therapeutic effects of PG-DFC-POP-PEP1 on inflammatory processes within a physiological context. (J) Pearson Coefficient Calculation from I for estimation of colocalization of IL 1β and NLRP3. Statistical comparisons were made using ANOVA test for equality of medians, with **, ***, and **** indicating significant differences (p < 0.05, p < 0.005, and p < 0.001, respectively), while NS denotes nonsignificant results. Data are presented as mean ± SEM.

## CONCLUSIONS

In this manuscript, we introduce a novel pH-dependent conformational switch-based peptide designed for selective cancer therapy through the Lysosome mediated Pyroptosis pathway. To overcome limitations in conventional cancer treatment, we synergistically combined two robust techniques: the OH-enriched PG-DGC-POP as an innovative drug carrier targeted specifically to cancer cells and the MPT63-based pH-dependent conformational switchable peptide, PEP1.Through comprehensive Biophysical and Simulation studies; we successfully validated the pH-dependent structural alterations of the peptide. Our investigations delved into the intricate N-GSDMD mediated Pyroptosis pathway, particularly via the Hsp70-LC3 axis, observed in both *in vitro* and *in vivo* Colon cancer models (Figure 7). Because of the PG-DFC-POP-PEP1 treatment, we observed lysosomal destabilizationthrough the LC3 axis.This phenomenon triggered an Inflammasome activated N-GSDMD mediated Pyroptosis pathway. We believe that the present study unveils a promising avenue for Peptide-based nanomedicine development, specifically targeting lysosomes for selective cancer treatment. Moreover, this systematic exploration of lysosome induced Pyroptosis will serve as a foundational resource for biologists, offering insights into diverse cell death mechanisms associated with cancer treatment.

**Figure 7:**
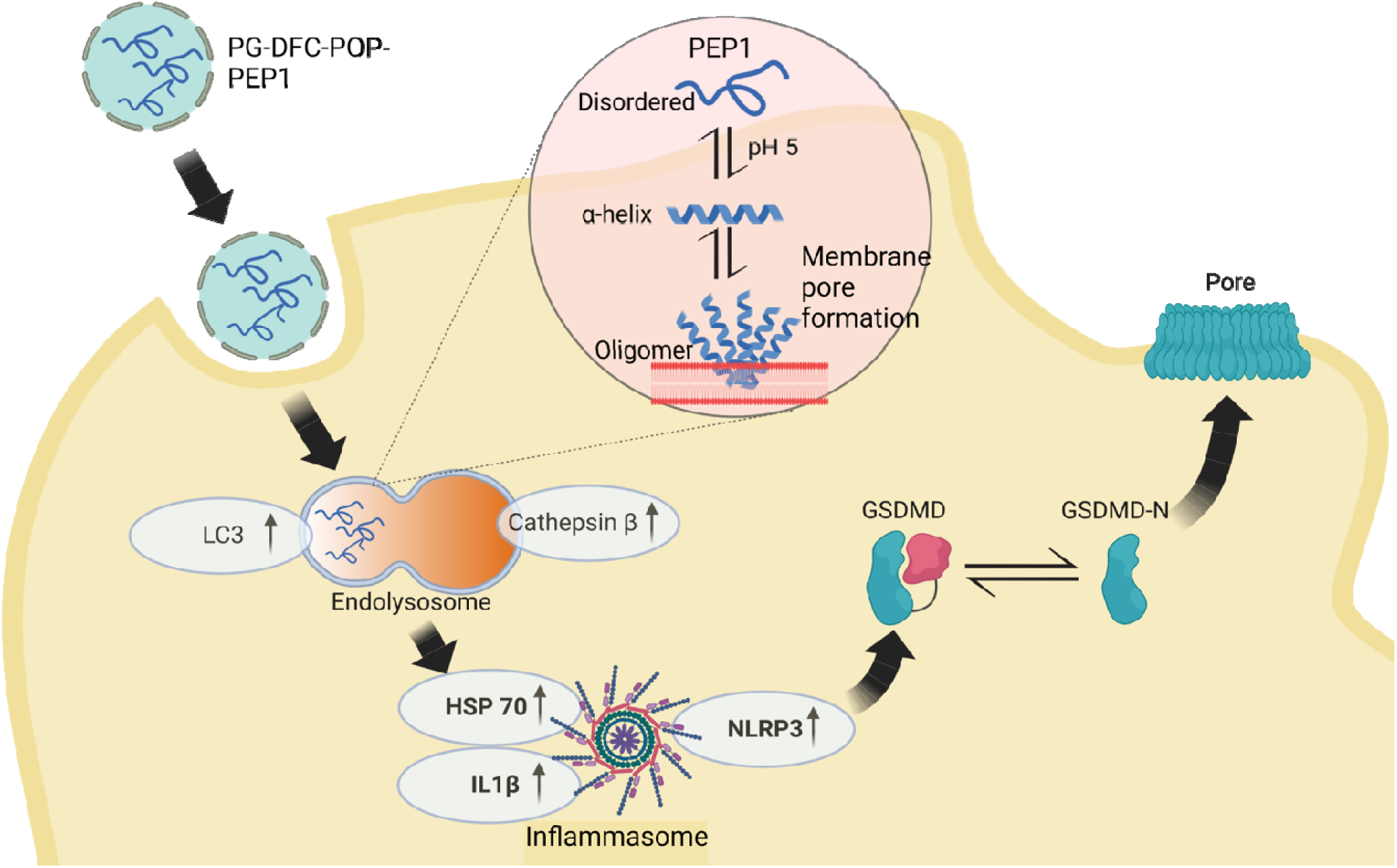
The schematic depiction illustrates the mechanism of cancer cell death via t**he** Pyroptosis pathway facilitated by PEP1 encapsulated within PG-DFC-POP nanomateri**al,** employing a conformational switch strategy activated by acidic pH conditions. The PG-DFC**-** POP nanocarrier, rich in multi-OH groups, selectively transports PEP1 to cancer cells. Followi**ng** cellular uptake, PEP1 is liberated from the PG-DFC-POP carrier within the acidic milieu of t**he** endolysosome. Upon release, PEP1 undergoes a conformational shift, adopting an alpha-heli**cal** structure, which then assembles into proteolysis-resistant oligomers. These oligomers proceed **to** perforate the endolysosomal membrane, inducing its destabilization. Subsequently, the Hsp70**-** LC3 axis is activated, culminating in heightened levels of IL1β, NLRP3, and cathepsin β, alongside inflammasome assembly. These molecular perturbations collectively instigate N**-** GSDMD-mediated programmed cell death in the cancer cell population.

## EXPERIMENTAL SECTION

### MATERIALS

The HCT-116 (human colorectal carcinoma), CCD 841CoN (Colon epithelial-like cell), HEK293 (Human embryonic kidney cell) cell lines were acquired from NCCS, Pune, India. Gibco-Life Technologies in Grand Island, NY, USA provided Dulbecco’s Modified Eagle’s Medium (DMEM), penicillin/streptomycin/neomycin (PSN) antibiotic, fetal bovine serum (FBS), trypsin, and EDTA. NUNC in Roskidle, Denmark, from Fermentas, EU provided tissue culture plastic wares. SRL in India, Invitrogen and Sigma-Aldrich from US provided 3-(4,5-Dimethylthiazol-2-yl)-2,5-diphenyltetrazolium bromide (MTT), DAPI (4’,6-diamidino-2-phenylindole dihydrochloride), acridine orange, and ethidium bromide, respectively. Antibodies were purchased from Santa Cruz Biotechnology, Inc. USA and eBioscience, Inc. San Diego, USA. Peptides were purchased from GenScript Biotech Corporation in New Jersey, U.S. Millipore water was utilized for all studies. All other chemicals were acquired from Sigma-Aldrich Co. in St Louis,USA. Alexa fluor-488 maleimide and calcein dyes were obtained from Invitrogen (Eugene, Oregon,USA). 1,2-Dioleoyl-sn-glycero-3-phosphocholine (DOPC,18:1), 1,2-dioleoyl-sn-glycero-3 -phosphoethanolamine(DOPE, 18:1), 1,2-dioleoyl-sn-glycero-3 -phospho-( 1 ’-rac-glycerol)(DOPG18:1), 1,2-dioleoyl-sn-glycero-3-phospho-L-serine (DOPS,18:1),Sphingosylphosphorylcholine (18:1), 1,2-dioleoyl-sn-glycero-3-phosphate were purchased from Avanti Polar Lipids Inc. (Alabaster, AL,USA).

## METHODS

### Synthesis of diformylcresol (DFC)

In a typical synthetic procedure^34^, initially p-cresol (1.08 g, 0.01 mol) was taken in 100 ml RB with 5 mL of acetic acid. Then, hexamethylenetetramine (0.02 mol, 2.82 g) and paraformaldehyde (0.1 mol, 3 g) were added to the reaction mixture. After that the reaction mixture was allowed to stir for 2 h at 80°C to obtain a light-brown viscous mixture. One mLH_2_SO_4_ was added dropwise into it after cooling down the mixture to room temperature, which was followed by refluxing for 30 min. Then, 40 mL of distilled water was added into the resulting solution and a pale-yellow precipitate appeared instantly. The reaction mixture was kept in a refrigerator overnight to enhance the product yield. Then, the product was filtered off and washed with water, followed by a small amount of methanol. Finally, the product was recrystallized from toluene to obtain the pure product.

### Synthesis of PG-DFC-POP

Phluroglucinol (2 mmol) and diformyl cresol (3 mmol) were taken in a 50 mL autoclave then 10 mL of 1, 4-dioxane was added into it and stirred for 10 minutes. After that autoclave was kept in a furnace of 220° C for 96 hours. Then a precipitate was obtained which was washed with dry methanol, acetone, and THF and dried to get a deep red color powder material.

### Computational Characterization of PEP1 by molecular dynamics simulation

To verify the conformational changes within the chameleon sequence stretches (PEP1) present in MPT63^35, 36^ classical atomistic molecular dynamics simulations have been performed with the help of the NAMD program. Initially, to mimic the experimental conditions, two (pH7 and pH2) input models of PEP1 have been constructed. For verification of random coil to helix transformation, protonation states of the titratable residues (VAL, LYS, ASP, TYR, CYS) were changed based^38^ on the pKa/pH correlation of the amino acids via CHARMM-GUI input generator. A peptide (PEP1) in a water box model was constructed. Na+and Cl^-^ ions were added as per the final salt concentration of 150mM, to ensure charge neutrality of the system. Molecular interactions were defined by the CHARMM36force field^38^. Within a 12 Å cut-off, the van der Waals and short-range electrostatic interactions were calculated, and the particle mesh Ewald (PME) technique was used to assess long-range electrostatic interactions^39^. Langevin dynamics and the Langevin piston algorithm were employed, to keep the temperature and pressure at 300 K and 1 atm, respectively. To investigate the strand to helix propensity, the PEP1 chameleon stretches were subjected to the mentioned condition for a 50 ns simulation trajectory.

### Characterizations of PG-DFC-POP

To obtain powder X-ray diffraction patterns of PG-DFC-POP samples, a Bruker D8 Advance SWAX diffractometer was utilized with a fixed voltage (40 kV) and current (40 mA). The XRD machine was calibrated using a conventional silicon source and Ni-filtered Cu Kα radiation using a wavelength of 0.15406 nm. The surface area analyzer QuantachromeAutosorb 1-C was used for nitrogen adsorption/desorption analysis at 77 K. Before gas adsorption, the sample was degassed for 12 h at 403 K under a high vacuum. The NLDFT pore size distributions were obtained using the carbon/cylindrical pore model from the N_2_ sorption isotherm.^40^ For theTEM analysis, 10 mg of both PG-DFC-POP and PG-DFC-POP-PEP1 samples were dispersed in absolute ethanol for 5 min under sonication, followed by sample coating on a copper grid and air-drying before analysis. JEOL JEM 6700 field emission-scanning electron microscope (FE SEM) was used to analyze the particle size and morphology of PG-DFC-POP samples. High-throughput biophysical spectral analysis of the samples was conducted, including FT-IR analysis using a Nicolet MAGNA-FT IR 750 spectrometer Series II and UV-visible diffuse reflectance spectroscopy (UV 2401PC).

### Peptide synthesis

^19^VGQVVLGWKVSDLKSSTAVIPGYC^40^(PEP1) and^52^ATVNAIRGSVTP AVSQFNARTADGC^74^(PEP2) were synthesized and supplied by Abgenex (KRIC distribut**or,** USA). Peptides are ∼ 95% pure.

### Fluorescence Correlation Spectroscopy (FCS) Experiments and Data Analysis

FCS experiments were carried out on Alexa 488 maleimide labeled PEP1 using a dual channel I**SS** Alba V system equipped with a 60X water-immersion objective (NA 1.2)^41^. The labeled peptide samples were excited using an argon laser at 488 nm. The values of τ_D_ obtained with the peptide samples were normalized using the τ_D_ value obtained with the free dye (Alexa488Maleimide) which was measured under identical conditions. This procedure was needed to minimize the contribution of any small changes in the viscosity and refractive indices.

For a single-component system, diffusion time (τ_D_) of a fluorophore and the average number **of** particles (N) in the observation volume can be calculated by fitting the correlation functi**on** [G<τ)] to:

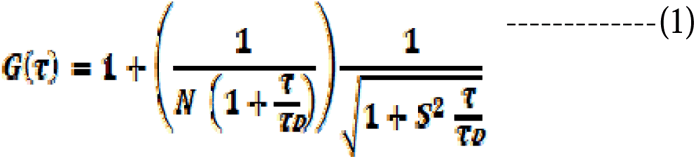

where, S is the structure parameter, which is the depth-to-diameter ratio. The characteris**tic** diffusion coefficient (D) of the molecule can be calculated from τ_D_ using:

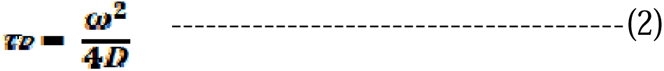

where, ω is the radius of the observation volume, which can be obtained by measuring the τ_D_**of** a fluorophore with known D value. The value of hydrodynamic radius (rH) of a labell**ed** molecule can be calculated from D using the Stokes-Einstein equation:

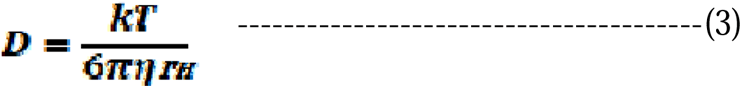

where, k is the Boltzmann constant, T is the temperature and η corresponds to the viscosity **of** the solution^42^.

### AFM studies

A 5 µL portion was taken from the diluted PEP1 sample and deposited on the freshly cleaved mica for 10 min. The typical peptide concentration was 500nM. After removi**ng** the excess liquid, the peptide samples were rinsed with MilliQ water and then dried using **a** stream of nitrogen. AFMimages were acquired at ambient temperature using a Bioscope Catal**yst** AFM (Bruker Corporation, Billerica, MA) with silicon probes. The standard tapping mode w**as** used to image the morphology of oligomers. The nominal spring constant of the cantilever w**as** kept at 20– 80 N/m. The spring constant was calibrated by a thermal tuning method. A standa**rd** scan rate of 0.5 Hz with 512 samples per line was used for imaging the samples. A single third**-** order flattening of height images with a low pass filter was done followed by section analysis **to** determine the dimensions of oligomers.

### Preparation of small unilamellar vesicles (SUVs)

An appropriate amount of DOPC lipid (other lipids like DOPS, DOPE, DOPG, SM, DOPA) **in** chloroform (concentration of the stock solution is 25 mg mL-1) was transferred to a 10 mL glass bottle. The organic solvent was removed by gently passing dry nitrogen gas. The sample was then placed in a desiccator, which was connected to a vacuum pump for about 2 hours to remove traces of the leftover solvent. A required volume of 20 mM sodium phosphate buffer at pH 7.4 and pH 5 was added to the dried lipid film so that the final desired concentration (10 mM) was obtained. The lipid-film with the buffer was kept overnight at 4°C to ensure efficient hydration of the phospholipid heads. Vortexing of the hydrated lipid film for about 30 min produced multilamellar vesicles (MLVs). Long vortexing was occasionally required to make uniform lipid mixtures. This MLV was used for optical clearance assay. For preparing the small unilamellar vesicles, MLVs were sonicated using a probe sonicator at amplitude 45% for 30 minutes and after that sample was centrifuged at 5000 rpm to sediment the tungsten artifacts and finally it was filtered by 0.22µm filter unit. Size of the small unilamellar vesicles was measured by DLS and the average diameter was found to be ∼ 70 nm.

### Circular Dichroism Measurement

Near and Far-UV CD spectra of PEP1atpH 7.5 and pH5 were recorded using a JASCO J720 spectro-polarimeter (Japan Spectroscopic Ltd.). Far-UV CD measurements (between 200 and 250 nm) were performed using a cuvette of 1 mm path length. A peptide concentration of 5mg/ml was used for the CD measurements. The scan speed was 50 nm min-1, with a response time of 2 s. The bandwidth was set at 1 nm. Three CD spectra were recorded in continuous mode and averaged.

### Assay for Permeabilization of Lipid Vesicles

The ability of peptides to promote the release of calcein from entrapped SUVs composed of different lipids was checked by monitoring the increase in fluorescence intensity of calcein. Calcein-loaded liposomes (70 mM concentration) were separated from non-encapsulated (free) calcein by gel filtration on a Sephadex G-75 column (Sigma) using an elution buffer of 10 mM MOPS, 150 mM NaCl and 5 mM EDTA (pH 7.4), and lipid concentrations were estimated by complexation with ammonium ferro-thiocyanate Fluorescence intensity was measured at room temperature (25°C) using a PTI spectro-fluorometer.A path length of 1cm was used for each measurement. The excitation wavelength was 490 nm and emission were set at 520 nm. Excitation and emission slits with a nominal bandpass of 3 and 5 nm were used, respectively. The high concentration (70 mM) of the entrapped calcein led to self-quenching of its fluorescence resulting in low fluorescence intensity of the vesicles (I_B_). Release of calcein caused by the addition of peptide led to the dilution of the dye into the medium, which could therefore be monitored by an enhancement of fluorescence intensity (If). This enhancement of fluorescence is a measure of the extent of vesicle permeabilization. The experiments were normalized relative to the total fluorescence intensity (It) corresponding to the total release of calcein after complete disruption of all the vesicles **by addition of Triton X-100 (2% v/v). The percentage of calcein release in the presence of peptides** was calculated using the equation:

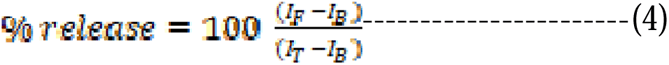

where, I_B_ is the background (self-quenched) intensity of calcein encapsulated in vesicles, I_F_ represents the enhanced fluorescence intensity resulting from the dilution of dye in the medium caused by the peptide induced release of entrapped calcein. I_T_ is the total fluorescence intensity after complete permeabilization is achieved upon addition of Triton X-100. Typical SU**Vs** concentration was taken 100 µM and 1µM concentration of each peptide sample was used.

### Cell Culture

CCD 841CoN, HEK293 and HCT116cells were grown in a medium known **as** Dulbecco’s Modified Eagle Medium (DMEM), which contains nutrients and suppleme**nts** required for cell growth. The medium was supplemented with 10% fetal bovine serum (FB**S),** which provided essential growth factors and nutrients for the cells. Additionally, 1% antibio**tic** cocktail was added to the medium to prevent contamination. The cells were incubated in **a** controlled environment, with a temperature of 37°C and a constant level of 5% carbon dioxi**de** (CO2), which helps to maintain physiological conditions. This environment was also humidified to prevent the cells from drying out. When the cells reached 75-80% confluence, meaning they had grown to cover 75-80% of the surface area of the culture vessel, they were harvested using a mixture of trypsin (0.25%) and EDTA (0.52 mM) in phosphate-buffered saline (PBS). Trypsin is an enzyme that cleaves proteins, which loosens the cells from the surface of the culture vessel, while EDTA was added to chelate the calcium ions, which aids in detaching the cells.

### MTT assay

To evaluate cell viability, an MTT [(4,5-dimethyl-thiazol-2-yl)-2,5-diphenyl tetrazolium bromide] assay^43^ was performed to assess the cytotoxic effects of PG-DFC-POP, PEP1andPG-DFC-POP-PEP1. Cells were seeded in 96-well plates at a density of 2 x 10^4^cells per well and treated with varying concentrations of PG-DFC-POP, PEP1, and PG-DFC-POP-PEP1.

The plates were then incubated at 37°C in a humidified environment containing 5% CO_2_ for 24 hours. After incubation, the cells were washed with phosphate-buffered saline (PBS), and MTT solution (4 mg/mL) was added to each well. The plates were again incubated for 4 hours, allowing the MTT to be converted to an intracellular formazan salt. The formazan salt was solubilized in dimethyl sulfoxide (DMSO), and the absorbance was measured at 595 nm using an ELISA reader (Emax, Molecular Device, USA). To ensure consistency; PG-DFC-POP and PG-DFC-POP-PEP1 samples were sonicated before treatment to obtain homogenized mixtures. The reported data represents the average of triplicate.

### Assessment of Pyroptotic Cell death under Optical Microscope

The morphology of HCT-116 cells was monitored using optical microscopy before and after treating them with 5 µg/mL of PG-DFC-POP-PEP1 for 4 and 8 h. Following treatment, the cells were incubated under standard conditions of 37 °C and 5% CO2 and then examined using a confocal laser scanning microscope (ZEISS LSM-980, Germany).

### Determination of cell death via Annexin-V staining

Overnight, HCT-116 cells were plated at a density of 7 × 10^5^ cells per well in a 6-well plate. The medium was replaced with fresh medium the next day, followed by treating the cells with 5 µg/mL nanoparticles for 4 and 8 hours, respectively. After 4 and 8 hours of incubation, cell pellets were collected and washed in PBS. Subsequently, the Annexin V-FITC Apoptosis Detection Kit was used to stain the cells according to the manufacturer’s instructions^44^. The stained cells were analyzed using a BD LSRFortessa flow cytometer (San Jose, CA, USA) by detecting Annexin V-FITC and PI staining in the FITC and PE channels, respectively. Finally, FlowJo software was used to analyze the flow cytometry data, identifying Annexin V/PI-positive cells as apoptotic populations and Annexin V-negative/PI-positive cells as populations undergoing non-apoptotic cell death.

### Animal ethics and maintenance

Five-week-old male C57BL/6 mice weighing between 14-16 grams were housed in optimal conditions with unlimited access to standard pellet food (comprised of 65.5% carbohydrate, 21% protein, 5.5% fat, 7% mineral mixture, and 1% vitamin mixture) and filtered water. The mice were placed in cages containing no more than five animals, in a controlled environment with a temperature range of 21-22°C, humidity maintained between 55-65%, and a 12-hour light/dark cycle. All animal experiments conducted adhered to guidelines set forth by the Institutional Animal Ethics Committee (IAEC) and the Committee for the Purpose of Control and Supervision of Experiments on Animals (CPCSEA), as prescribed by the Government of India.

### *In vivo*Experimentation

On the first day, the animals were shaved on the back flank. In the shaved right flank of each mouse, B16F10 (2 × 10^6^) in PBS was injected subcutaneously. Eight days after tumor implantation, the animals were randomly divided into four groups as follows (five mice in each group): group I, served as a control and mice in this group were kept at a standard ambient temperature of 24 ± 2 °C and 60-70% relative humidity; group II, B16F10 tumor-bearing mice treated with normal saline; groups III and IV, B16F10 tumor-bearing mice treated with PG-DFC-POP-PEP1 (1 and 1.5 mg/kg bodyweight) intra-peritoneally for six alternate days. The resultant tumor length and weight were measured after the completion of the experimentation. In addition, the animal survival rate was evaluated up to 30 days.

## ELISA

To prepare the samples, the Bullet Blender Storm was used for mechanistic homogenization. Total proteins were extracted from the samples using RIPA lysis buffer (Sigma Aldrich, MO), which included a protease inhibitor, phosphatase inhibitor (Roche, Basel, Switzerland), and PMSF (Sigma Aldrich, MO). After centrifugation at 12,000 rpm for 15 minutes at 4°C, the resulting supernatants containing the total protein were subjected to mHGF ELISA (R&D systems, MN), following the manufacturer’s instructions^45^.

### Immunocytochemistry

HCT-116 cells were cultured on cover slips for an initial period of 24 hours. Following this, the cells were subjected to treatment with Alexa 488 conjugated PG-DFC-POP-PEP1 at a concentration of 5 µg/mL, and subsequently incubated for 4 and 8 hours. Control cells that were not subjected to treatment were also included. The control and treated cells were then subjected to two washes with PBS (0.01 M) lasting 10 minutes each, and subsequently blocked for a period of 1 hour with a solution containing 0.3% Triton X-100 and 2% normal bovine serum in PBS. Next, the cells underwent an overnight incubation at a temperature of 4 °C with the appropriate primary antibody. Following this incubation, the cells were washed and subsequently incubated with the appropriate fluorophore-conjugated secondary antibodies for a duration of 2 hours. In certain experiments, the addition of Lysotracker Red was also employed. The cells were finally counterstained with DAPI and mounted with the Prolong Anti-fade Reagent (Molecular Probe, Eugene, OR), before being visualized via confocal laser-scanning microscopy (FV 10i, Olympus, Japan)^46^.

### Western Blotting

To prepare the tissue or cell lysate, a lysis buffer was utilized that contained Tris–HCl (20 mM) with a pH of 7.5, 2-mercaptoethanol (50 mM), EGTA (5 mM), EDTA (2 mM), NP40 (1%), SDS (0.1%), deoxycholic acid (0.5%), NaF (10 mM), PMSF (1 mM), leupeptin (25 mg mL-1) and aprotinin (2 mg mL-1). The protein concentration was determined using a BCA assay protocol, and the protein fractions were reconstituted in a sample buffer containing Tris–HCl (62 mM), SDS (2%), glycerol (10%), and β-mercaptoethanol (5%). The protein samples were denatured by boiling at 97 °C for 6 minutes. Next, equal amounts of denatured protein samples (25 µg) were separated using SDS-PAGE on a polyacrylamide gel (12.5%) and transferred overnight onto polyvinylidene difluoride membranes (PVDF). The PVDF membranes were then blocked with a mixture of non-fat dried milk (5%) in TBS containing Tween-20 (1%) for 1 hour at room temperature, followed by overnight incubation with primary antibodies (diluted at 1:1000). A second incubation was carried out with the secondary antibody coupled with either horseradish peroxidase (HRP)-conjugated goat anti-rabbit or anti-mouse antibodies (diluted at 1:3000) for 2 hours after washing the blots with TBS containing Tween-20 (1%). Finally, the blots were detected using an enhanced chemiluminescence (ECL) reagent, and the images were captured using a Chemidoc XR Imager (Bio-Rad Laboratories, Hercules, CA, USA)^47^.

### Immunohistochemistry

To perform histological analysis, the tumour region and adjacent region of the lung were collected, washed with PBS, and fixed with 4% paraformaldehyde in PBS for 2 hours at room temperature. Paraffin embedding was carried out by sectioning the tissue to a thickness of 5 µm, and standard Trichrome staining was performed using a staining kit from Merck-Sigma-Aldrich Corporation, St. Louis, Missouri, USA. The slides were analyzed under a light microscope (Model BX51 TRF, Olympus, Japan).For immunohistochemistry^48^, tissue sections were deparaffinized and treated with 0.3% hydrogen peroxide for 15 minutes, followed by incubation with RetrivergenA (pH 5.6) (BD Bioscience, Franklin Lakes, NJ). After blocking with 2.5% BSA for 90 minutes at 4°C, the sections were incubated with primary antibodies at a 1:100 dilution for 12 hours at 4°C. Subsequently, fluorescence-tagged secondary antibodies were applied at a 1:500 dilution for 4 hours. Counterstaining was done with DAPI, and slides were mounted with n-propylgallate containing 50% glycerol. Finally, the slides were visualized and analyzed using a Leica Confocal Microscope and LAS X Life Science™ software.

### Statistical Analysis

The results were expressed as the Mean ± SEM derived from multiple data points. Statistical significance in the observed differences was determined through analysis of variance (ANOVA) utilizing GraphPad prism (9.0) software. A significance level of p < 0.05 was considered indicative of meaningful differences.

## Supporting information

Additional figures S1-S5.

## Acknowledgement

AB would like to thank DST-SERB, New Delhi for a Core Research Grant (Project Number: CRG/2022/002812).

