## Supplemental figures S1-S5 for "Conformational switch of a peptide provides a novel strategy to design peptide loaded porous organic polymer for Pyroptosis pathway mediated cancer therapy"

**^§^ Dept. of Cell, Developmental, & Integrative Biology, University of Alabama, Birmingham, AL 35233**

**^‡^Molecular genetics department, University of Texas Southwestern Medical center, USA**

^||^ **NIPM and SoLs, University of Nevada Las Vegas, Nevada, USA**

**^††^ School of Materials Science and Nanotechnology, Jadavpur University, Kolkata - 700032, India**

**^%^Department of Biology, New Mexico State University, Las Cruces, NM, USA**

**^‡‡^ Cancer Biology and Inflammatory Disorder Division, CSIR-Indian Institute of Chemical Biology, Kolkata, India**

**^#^School of Materials Sciences, Indian Association for the Cultivation of Science, Jadavpur, Kolkata 700 032, India**

**^⊥^ Structural Biology and Bioinformatics Division, CSIR-Indian Institute of Chemical Biology, Kolkata, India**

**^$^Department of Physics, University of Illinois at Urbana-Champaign, Urbana, United States**

**^&^Department of Chemical and Biomolecular Engineering, University of Illinois Urbana-Champaign, Urbana, 1801, Illinois, USA**

**Email addresses of the corresponding authors:**

****

**Figure S1:**Conformational switch in PEP1.(A).Plot of residue wise hydrophobicity of PEP 1 and PEP 2 as analysed by ProtScale (<http://web.expasy.org/protscale/>). (B). Aggregation Hotspot regions were identified using AGGRESCAN analysis. (C).Molecular dynamics simulation derived trajectories shows the residue specific conformational changes of PEP1 at pH (i) 7 and (ii) low pH. Here, different colors indicate different conformations. Leisure blue: loops and turns,yellow:extended conformation, blue:3-10 helix, pink: α helix, white: coil(D). Plot of RMSD with time for PEP1 at pH7 and at low pH as obtained from molecular dynamics simulation data.(E).AFM topographic image of PEP 1 at pH 5. Scale bar is 200 nm.

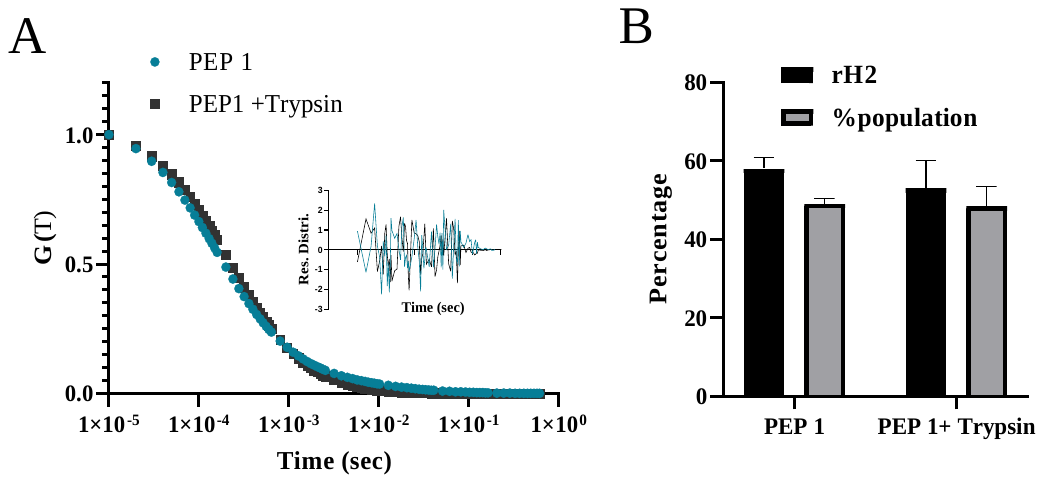

**Figure S2:** PEP1 oligomers are proteolysis resistant. (A). Correlation spectra of PEP 1 at pH5 in presence and absence of trypsin. Inset shows the residual distribution of the fit of the correlation function data. The randomness of the residual distribution stands for the randomness of the fit. (B). Bar plot of the hydrodynamic radii and percentage population of the respective particles (slow components that correspond to rH2, the hydrodynamic radii of the oligomeric species of PEP 1 at pH 5) in presence and absence of trypsin protease(PEP1 was incubated at pH5 with trypsin at 1:200 enzyme:substrate weight ratio at 24 °C).

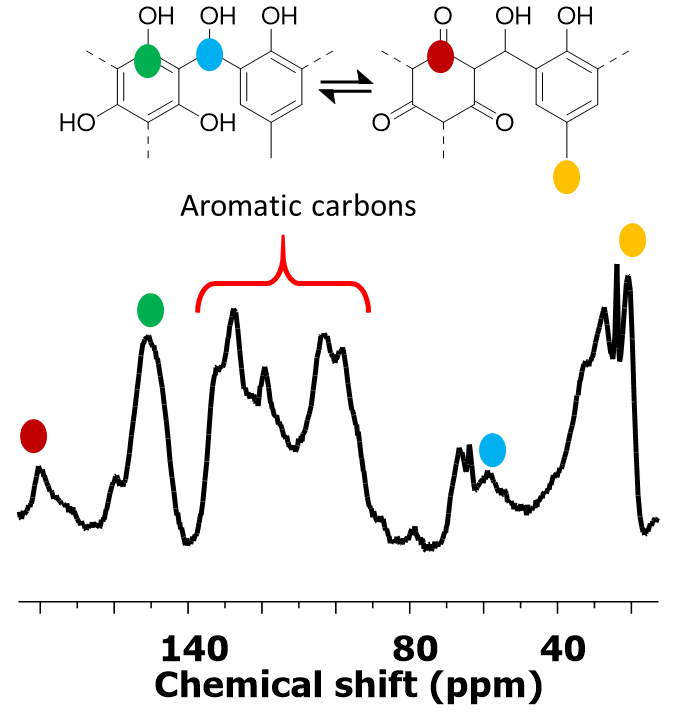

**Figure S3**. ^13^C solid NMR spectra of PG-DFC-POP.

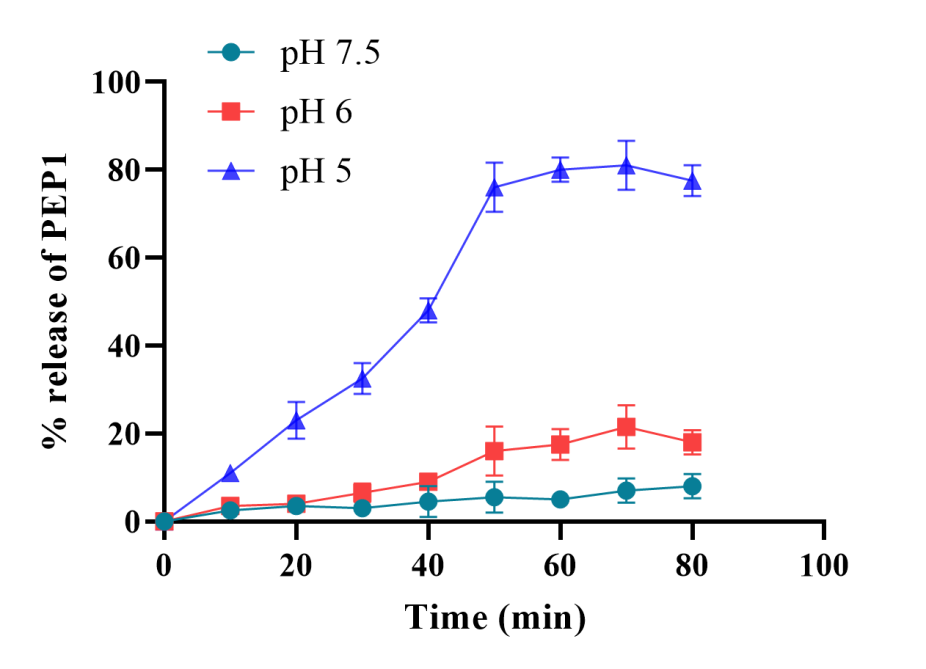

**Figure S4:**Release of PEP1 fromPG DFC POP PEP1. Plot shows the percentage of release of PEP 1 from PG DFC POP PEP1 nanomaterials with time at different pH conditions.

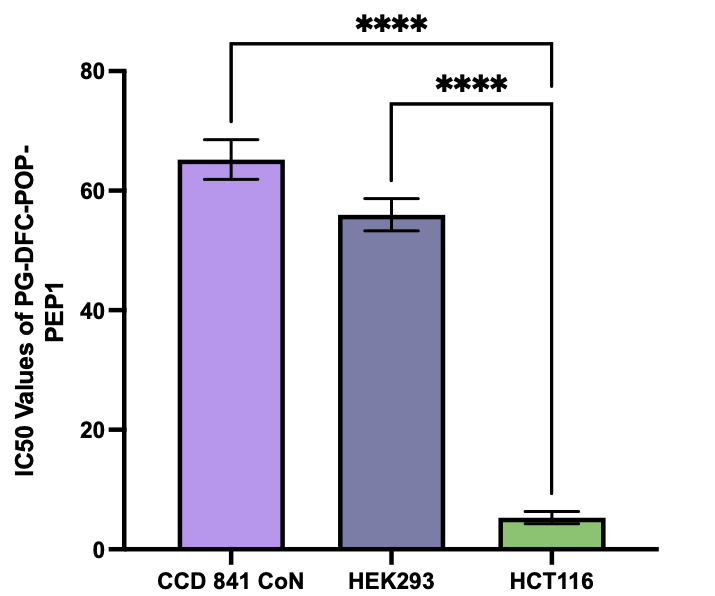

**Figure S5:** Cellular Cytotoxicity by PG-DFC-POP-PEP1 treatment in CCD 841CoN, HEK293 and HCT116 cell line. Here, IC50 values are expressed as μg/mL.
